# A Whole-Brain Regression Method to Identify Individual and Group Variations in Functional Connectivity

**DOI:** 10.1101/2020.01.16.909580

**Authors:** Yi Zhao, Brian S. Caffo, Bingkai Wang, Chiang-shan R. Li, Xi Luo

## Abstract

Resting-state functional connectivity is an important and widely used measure of individual and group differences. These differences are typically attributed to various demographic and/or clinical factors. Yet, extant statistical methods are limited to linking covariates with variations in functional connectivity across subjects, especially at the voxel-wise level of the whole brain. This paper introduces a generalized linear model method that regresses whole-brain functional connectivity on covariates. Our approach builds on two methodological components. We first employ whole-brain group ICA to reduce the dimensionality of functional connectivity matrices, and then search for matrix variations associated with covariates using covariate assisted principal regression, a recently introduced covariance matrix regression method. We demonstrate the efficacy of this approach using a resting-state fMRI dataset of a medium-sized cohort of subjects obtained from the Human Connectome Project. The results show that the approach enjoys improved statistical power in detecting interaction effects of sex and alcohol on whole-brain functional connectivity, and in identifying the brain areas contributing significantly to the covariate-related differences in functional connectivity.

## 1 Introduction

In experiments of resting-state functional magnetic resonance imaging (fMRI), the study of connectivity to characterize cerebral functional segregation and functional integration has received considerable attention. The understanding of brain functional organization may provide critical insights to cognitive function, as well as mental diseases. Functional connectivity, defined as the correlation or covariance between fMRI time courses, reveals the level of synchrony in the fluctuations of blood oxygenation level dependent (BOLD) signals between brain regions (Friston, 1994). Brain regions with high functional connectivity are generally grouped as functionally related and defined as a functional module/subnetwork. For example, the default mode network (DMN) is a functional subnetwork that shows greater activity during resting states than during many task challenges, and has been consistently identified through resting-state functional connectivity analysis (Greicius et al., 2003). Existing literature has shown that brain functional connectivity varies with respect to individual characteristics, such as sex and age (Scheinost et al., 2015; Lopez-Larson et al., 2011), and in patients with autism spectrum disorders (Assaf et al., 2010), Alzheimer’s disease (Wang et al., 2007), schizophrenia (Lynall et al., 2010) and other psychiatric disorders, as compared to healthy controls.

To describe group-level differences in brain functional connectivity, investigators typically perform statistical analysis on each individual connection. One critical drawback of this element-wise approach is the issue of multiplicity. That is, with *p* brain voxels/regions, statistical inference needs to account for at least *p*(*p* − 1)*/*2 hypothesis tests, one for each matrix element. To circumvent this, Zhao et al. (2019) proposed a whole-matrix regression approach called Covariate Assisted Principal (CAP) regression. It aims to identify a common linear projection of *p* time courses across subjects so that variations in functional connectivity defined by the projection can be explained by the covariates of interest. It is considered as a *meso*-scale approach in the sense that with an appropriate thresholding, the projection defines a brain subnetwork. However, this approach suffers from the so-called “curse of dimensionality”, in that the dimension of the data, *p*, cannot be greater than the number of fMRI volumes. Therefore, it cannot be directly applied to voxel-level fMRI data.

In fMRI studies, independent component analysis (ICA) is a widely used technique to cluster brain voxels into subnetworks (Beckmann, 2012). Applied to resting-state fMRI data, spatial ICA identifies spatially independent and temporally coherent components. Based on the biological assumptions regarding spatial contiguity of brain networks across individuals, group ICA was introduced for population-level studies (Calhoun et al., 2001). The time courses of all subjects are concatenated to identify independent spatial components shared by the study population.

In this study, we propose a method that integrates group ICA with CAP regression aiming to discover brain subnetworks that are associated with population/individual characteristics of interest. The method consists of three steps: (1) group ICA of the resting-state fMRI data; (2) CAP regression on the IC time courses; and (3) reconstruction of brain networks. The proposed method scales up the CAP regression in fMRI application to voxel-level analyses, and redefines brain subnetworks that demonstrate population/individual variation in functional connectivity. Here, we consider group ICA in the dimension reduction step as we aim to identify spatially independent brain networks that are shared across individuals. Other methods that identify common components in large scale data – group principal component analysis (PCA, Smith et al., 2014), for instance – can also be applied to this end.

This paper is organized as follows. In Section 2, we introduce our method. Section 3 presents an application in resting-state fMRI data obtained from the Human Connectome Project (HCP). Section 4 summarizes results with a discussion.

## 2 Method

Let 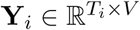 denote the *T*_*i*_ BOLD scans of *V* voxels acquired in the resting-state fMRI study from subject *i* (*i* = 1, …, *n, n* is the number of subjects). Let **x**_*i*_ ∈ ℝ^*q*^ be a covariate vector collected from subject *i*. The method aims to perform whole-brain functional connectivity analysis, to produce functional connectivity maps that vary with the covariates. Here, functional connectivity map means those voxels/areas that contribute largely to the covariate-related variations in functional connectivity across subjects.

We first implement group independent component analysis (ICA, Calhoun et al., 2001) to the whole brain data; then employ covariate assisted principal (CAP) regression (Zhao et al., 2019) to identify projections of the ICs that are associated with the covariates of interest; and finally we reconstruct the CAP brain maps. Figure 1 summarizes the data analysis pipeline. We elaborate on each step in the following sections.

**Figure 1:**
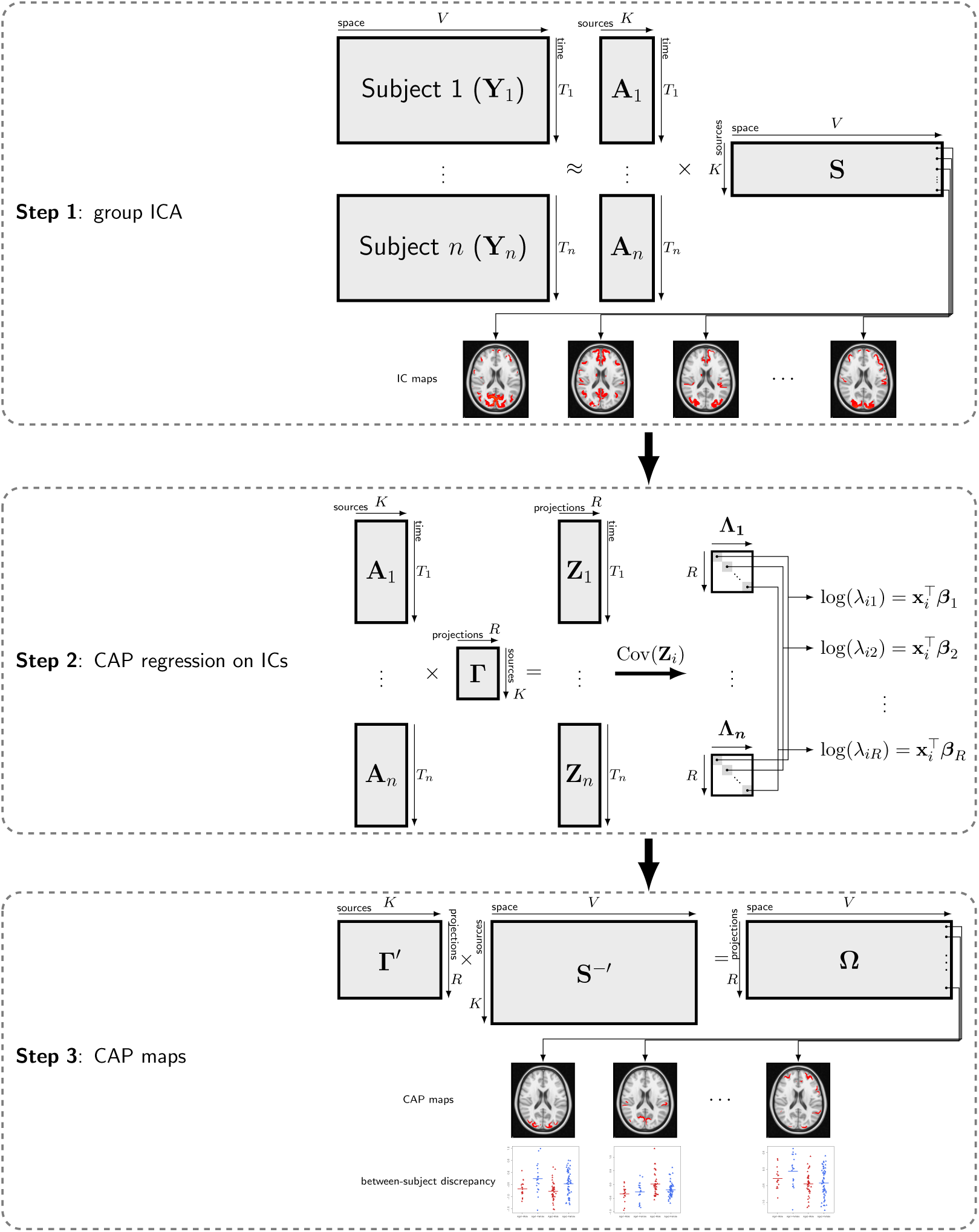
Data analysis pipeline. Step 1: group independent component analysis (ICA) on the whole brain. Step 2: the covariate assisted principal (CAP) regression on the IC time courses to identify projections of the ICs that are associated with the covariates of interest. Step 3: reconstruction of the CAP brain maps.

### 2.1 Group ICA

Independent component analysis was introduced to decompose multivariate data into a linear mixture of latent sources assuming that the latent components are non-Gaussian and statistically independent. Consider an *m*-dimensional random vector **x**, a noise-free ICA model assumes:

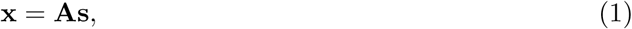

where **A** ∈ ℝ^*m*×*n*^ (*n* ≤ *m*) is a full rank mixing matrix and **s** ∈ ℝ^*n*^ is a *n*-dimensional independent random sources.

In fMRI studies, based on the sparse distributed nature of the spatial pattern for typical cognitive activation paradigms (McKeown et al., 2003), spatial ICA has been successfully implemented to discover spatially independent and temporally coherent brain regions. To elaborate, for BOLD signal **Y** ∈ ℝ^*T* ×*V*^ acquired from a single subject, spatial ICA assumes

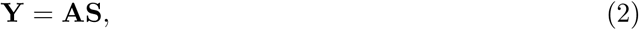

where **A** ∈ ℝ^*T* ×*K*^ is the scalar mixing matrix and **S** ∈ ℝ^*K*×*V*^ is the spatial independent component maps. It is assumed that the rows of **S** are statistically independent. However, it is not straight-forward to make population level inference based on individual level IC decompositions. Calhoun et al. (2001) introduced group ICA for drawing group level inference by temporally concatenating BOLD time courses from multiple subjects. The method assumes common spatial brain maps across subjects (see Step 1 in Figure 1). In practice, in order to reduce the computation complexity and the amount of required memory, multiple data reduction steps, typically using principal component analysis (PCA), are performed before concatenating the time courses (Calhoun et al., 2009).

### 2.2 Covariate assisted principal regression

Covariate assisted principal regression aims to identify the optimal data rotation such that data variation can be best interpreted by subject characteristics of interest. In fMRI studies, covariance/correlation matrices are generally implemented to characterize the temporal coherence of BOLD signals, so-called brain functional connectivity (Friston, 2011). Given the IC time courses **A**_*i*_’s attained from group ICA in the first step, the CAP approach assumes that there exist *R* ∈ {1, …, *K*} common diagnolizations of **A**_*i*_’s covariance matrices and that data variation in the projection space satisfies a log-linear model of the covariates (denoted by **x**_*i*_ ∈ ℝ^*q*^). Let **A**_*it*_ = (*A*_*it*1_, …, *A*_*itK*_)^T^ denote the *t*th row of **A**_*i*_ (for *t* = 1, …, *T*_*i*_, where *T*_*i*_ is the number of fMRI volumes of subject *i*) and ***γ***_*r*_ a vector in ℝ^*K*^. For *r* = 1, …, *R*, the CAP approach assumes that

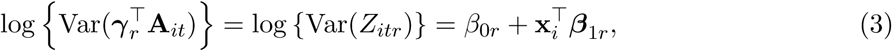

where 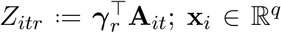 is the vector of covariates of interest; *β*_0_ and ***β***_1*r*_ ∈ ℝ^*q*^ are model coefficients; 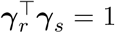 if *r* = *s* and 0 otherwise imposing orthogonality between projections. Zhao et al. (2019) proposed to estimate projection directions and model coefficients by maximizing the likelihood function assuming **A**_*it*_ is normally distributed with mean zero and covariance matrix **Σ**_*i*_ (for *t* = 1, …, *T*_*i*_ and *i* = 1, …, *n*). The number of projections, *R*, is determined based on the level of deviation from diagonality, as introduced in Zhao et al. (2019).

#### 2.2.1 Interpreting the CAP model under a special case

Though the CAP model aims to study global changes in functional connectivity, the regression coefficients have a simple interpretation under a special case. Assume ***γ***_*r*_ = (*γ*_*r*1_, …, *γ*_*rK*_)^T^ and **Σ**_*i*_ = (σ_*ijk*_)_*j,k*_ for *i* = 1, …, *n*, then

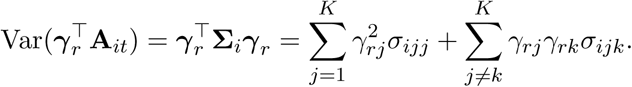

Consider the special case that only two entries in ***γ***_*r*_ are nonzero and identical. Without loss of generality, assume 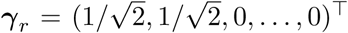, which indicates that this component consists of the first two ICs. Since the IC time courses are standardized to have homogeneous variance, for example all time courses are scaled with standard deviation one (that is, σ_*ijj*_ = 1 for ∀ *i* and ∀ *j*), then with σ_12_ = σ_21_,

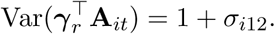

The regression model (3) assumes

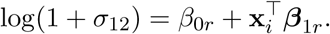

In the classic edge-wise regression model, it is assumed that the Fisher *z*-transformed Pearson correlation fits the linear regression model. The Fisher *z*-transformation takes the form

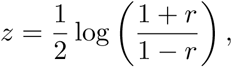

where *r* is the Pearson correlation between two ICs. When σ_*i*12_ ≈ 0, we have

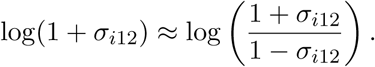

Therefore, under this special case, the CAP regression coefficients have a similar interpretation as those of edge-wise regression models.

### 2.3 Covariate related brain maps

The CAP procedure attains projections of the ICs. Integrating the spatial IC maps, we build new brain maps that explain association with the covariates in the terms of brain functional connectivity. Let **Γ** = (***γ***_1_, …, ***γ***_*R*_) ∈ ℝ^*K*×*R*^ denote the orthonormal projection matrix estimated from the CAP procedure. Each row of **Ω** = **Γ**^T^**S**^−T^ ∈ ℝ^*R*×*V*^ then gives the reconstructed CAP brain map (Step 3 in Figure 1), where **S**^−^ ∈ ℝ^*V* ×*K*^ is the generalized inverse of matrix **S**. Each CAP brain map, after threholding, should be interpreted as major brain areas contributing to specific functional connectivity variations explained by the covariates, especially those areas with statistically significant regression coefficients.

## 3 Analysis of resting-state fMRI from the Human Connectome Project

We apply our analysis pipeline to the Human Connectome Project (HCP) resting-state fMRI data. The HCP aims to characterize human brain structure, function and connectivity, as well as their variability in healthy adults. We use the group ICA data from the HCP, as available at http://www.humanconnectomeproject.org/. The fMRI data were first minimally-preprocessed following Glasser et al. (2013). The artifacts were removed by using ICA+FIX (Salimi-Khorshidi et al., 2014; Griffanti et al., 2014). Group-PCA results were first generated by MIGP (MELODIC’s Incremental Group-PCA) from 820 subjects, and then fed into group-ICA using FSL (https://fsl.fmrib.ox.ac.uk/fsl/fslwiki/FSL) MELODIC tool (Hyvarinen, 1999; Beckmann and Smith, 2004). Spatial-ICA was acquired in grayordinate space (surface vertices plus subcortical gray matter voxels, Glasser et al., 2013) at various dimensionalities.

In this study, we use the 25-IC data of 109 subjects (aged 22–36) from the HCP S500 release. The goal is to discover brain networks, within which the functional connectivity varies due to alcohol use, and to examine whether the alcohol-induced variation differs by sex. We apply the proposed method (i.e., ICA-CAP) and compare with an edge-wise regression approach. In both approaches, the regression model includes age (continuous, mean 29.0, SD 3.4), sex (binary, 41 females and 68 males), alcohol drinker (binary, 67 non-drinkers and 42 drinkers), and a sex×alcohol interaction (27 female non-drinkers, 40 male non-drinkers, 14 female drinkers and 28 male drinkers) as the covariates. In the following, we will focus on the four contrasts derived from the sex×alcohol interaction; that is (1) Male vs. Female among alcohol non-drinkers; (2) Male vs. Female among alcohol drinkers; (3) Alcohol drinkers vs. non-drinkers in the female group; and (4) Alcohol drinkers vs. non-drinkers in the male group.

In edge-wise regression, functional connectivity between IC’s is first calculated using the Pearson’s correlation and then Fisher *z*-transformed. Linear regression is performed with the Fisher *z*-transformed correlations as the outcome. We present the corresponding effect size in Figure 2 for those pairwise correlations that are significant for any contrast (at *α* = 0.05). We observe differences in functional connectivity between IC’s for all four comparisons. However, none of them survives correction for multiple testing following the false discovery rate control procedure in Benjamini and Hochberg (1995). Though the edge-wise regression approach identifies subtle variations in functional connectivity and the interpretation is straightforward, the method suffers from the curse of dimensionality as the number of tests increases dramatically as the number of components increases.

**Figure 2:**
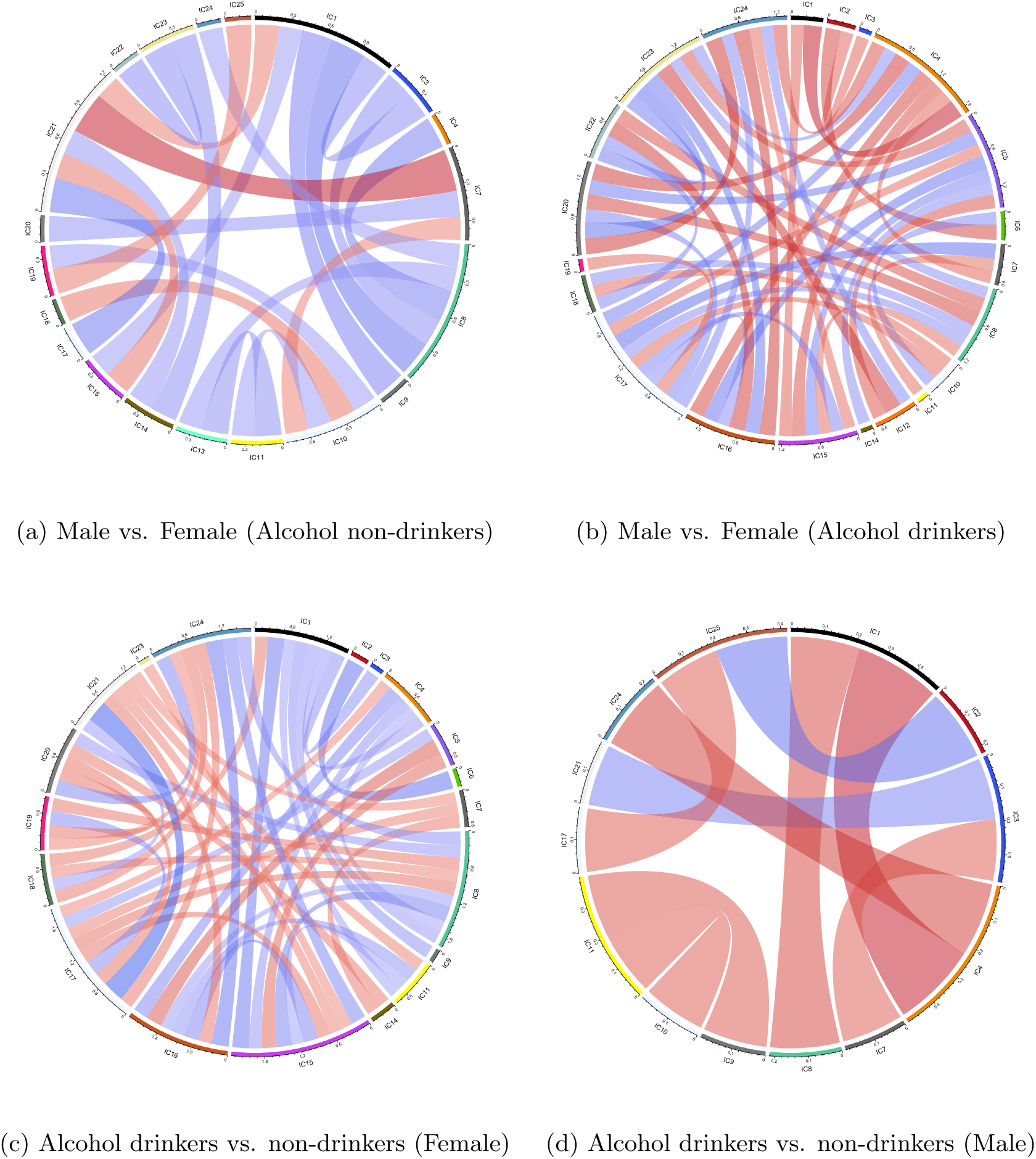
Effect size with significance of the model contrast of sex and alcohol in the edge-wise regression. The significance is evaluated if the raw *p*-value< 0.05. Red color indicates a positive effect and blue indicates negative. The darkness of the color and the width of the cord suggest the magnitude of the effect.

Using the deviation from diagonality to select model order, the proposed ICA-CAP approach identifies five components. Figure 3 shows the loading profile and Table 1 presents the coefficient estimates (with 95% bootstrap confidence intervals) of the five components. For the components C1, C2 and C4, we observe significant sex difference in functional connectivity among alcohol drinkers. In addition, among females, the functional connectivity within the component network demonstrates a significant difference between alcohol drinkers and non-drinkers. For C3, both the alcohol drinkers and non-drinkers groups show significant sex difference. For C5, all four comparisons reveal significant difference in functional connectivity. Figure A.1 in the Supplement presents the scatter plot of each sex×alcohol subgroup for the five components. For C1 and C3, we fit the edge-wise regression model on the two top loading ICs and compare the results with CAP in Figure 5. The significance of the *β* coefficients of the top loading pair in the element-wise regression is consistent with those in the CAP components, which verifies the ICA-CAP findings.

**Table 1:** Estimated model contrast (and 95% bootstrap confidence interval) of sex and alcohol for the five identified components from the ICA-CAP approach.

|  | Male vs. Female |  | Alcohol drinkers vs. non-drinkers |  |
| --- | --- | --- | --- | --- |
|  | Alcohol Non-drinkers | Alcohol drinkers | Female | Male |
| C1 | -0.026 (-0.287, 0.241) | -0.461 (-0.785, -0.129) | 0.530 ( 0.157, 0.882) | 0.095 (-0.131, 0.307) |
| C2 | -0.002 (-0.189, 0.202) | -0.421 (-0.624, -0.198) | 0.343 ( 0.093, 0.585) | -0.075 (-0.247, 0.089) |
| C3 | -0.359 (-0.579, -0.160) | -0.290 (-0.517, -0.067) | -0.209 (-0.469, 0.050) | -0.140 (-0.306, 0.020) |
| C4 | 0.014 (-0.191, 0.236) | 0.289 ( 0.010, 0.559) | -0.333 (-0.602, -0.079) | -0.058 (-0.290, 0.150) |
| C5 | 0.351 ( 0.203, 0.534) | -0.165 (-0.336, -0.008) | 0.326 ( 0.157, 0.496) | -0.190 (-0.368, -0.027) |

**Figure 3:**
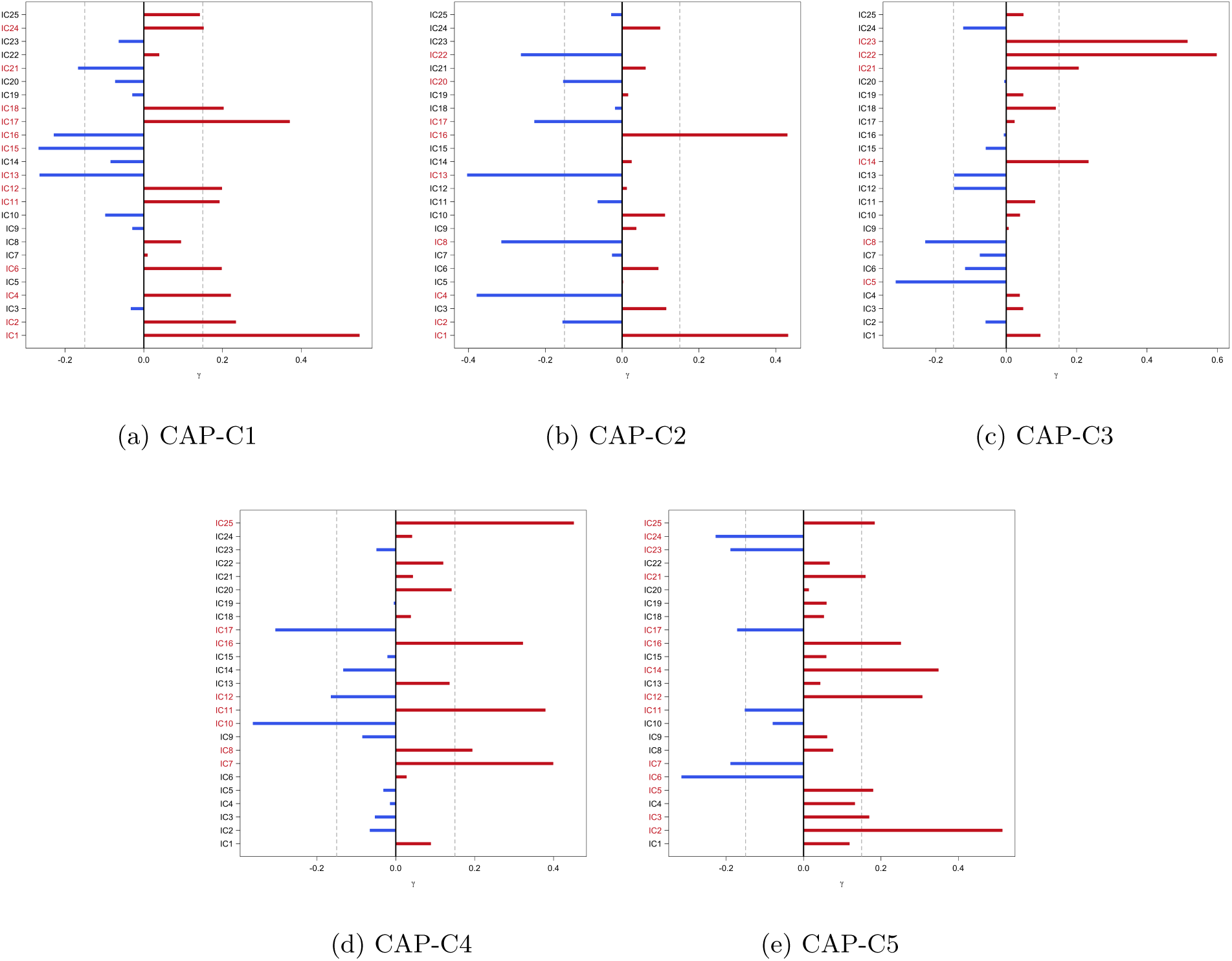
Loadings of the five identified components from the ICA-CAP approach. ICs in red have loading magnitude greater than 0.15 (gray dashed lines).

**Figure 5:**
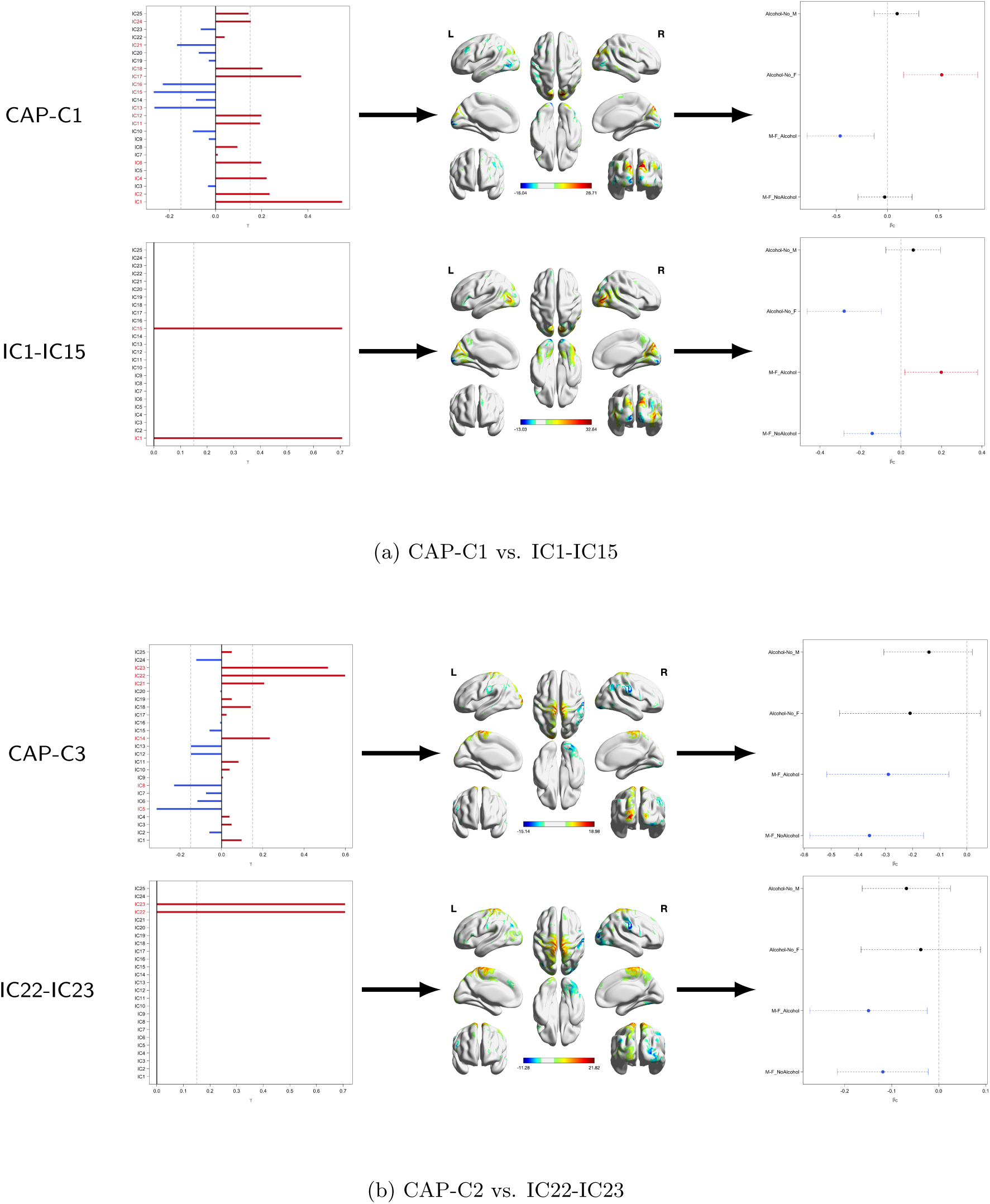
A comparison of CAP components with element-wise regression. Figures on the left panel shows the loading profile of the components/pairs, in the middle displays the corresponding brain map, and on the right presents the estimate of the four comparisons with 95% confidence intervals.

We highlight the regions with high loadings in the reconstrcuted CAP brain map in Figure 4. Of the five components, we use C1 and C3 as an example to further interpret the findings. For C1, the precuneus is among the highlighted regions. The precuneus is implicated in cue-elicited craving and altered emotion processing in alcohol dependence (Jansen et al., 2019). Individuals with alcohol use disorder demonstrated decreases in cortical thickness in a number of brain regions that include the precuneus (Schmidt et al., 2017). Compared with healthy controls, alcohol dependent participants showed lower degree centrality values in the cerebellum, visual cortex and precuneus in graph theoretical connectivity analyses (Luo et al., 2017). Another connectivity study reported that the precuneus, postcentral gyrus, insula and visual cortex were the main brain areas with reduction in network connectivity, perhaps suggesting reduced interoceptive awareness in alcohol drinkers, compared to non-drinkers (Vergara et al., 2017). Overall, the current findings are consistent with these earlier studies.

**Figure 4:**
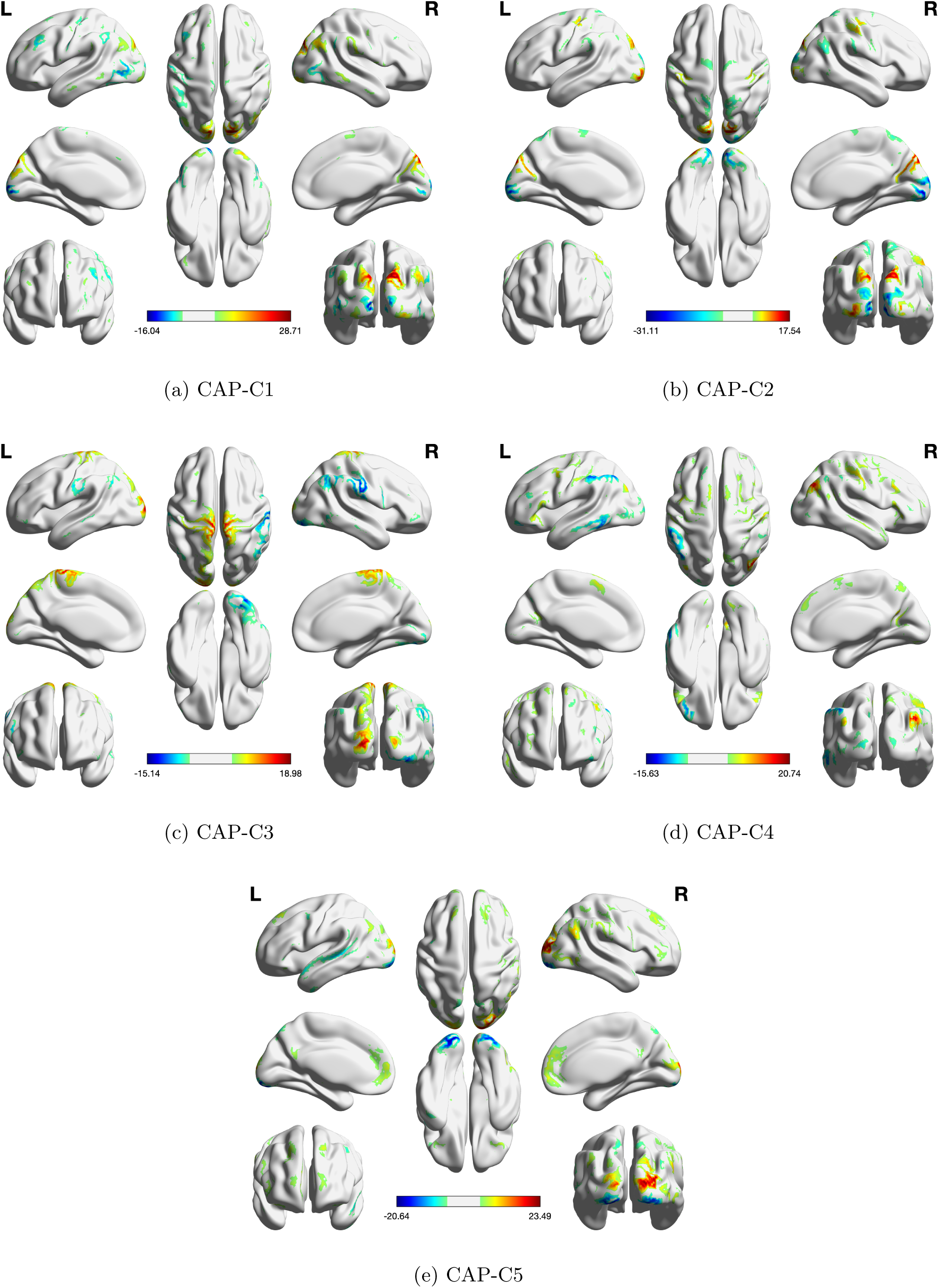
Reconstructed brain maps of the five components from the ICA-CAP approach.

For C3, we observe sex difference in brain functional connectivity. Very few studies have examined sex differences in resting state functional connectivity in neurotypical populations within the age range of the current cohort. An earlier study employed ICA to identify four fronto-parietal networks and showed sex differences in two of these networks with women exhibiting higher functional connectivity in general, an effect that appeared to be independent of the menstrual cycle (Hjelmervik et al., 2014). A life span study showed that the differences in connectivity between men and women of 22–25 years of age did not differ significantly in functional connectivities (Conrin et al., 2018). However, the 26–30 (*p* = 0.003) and the 31–35 age groups (*p* < 0.001) showed significant differences. At the most global level, areas of diverging sex difference include parts of the prefrontal cortex and the temporal lobe, amygdala, hippocampus, inferior parietal lobule, posterior cingulate, and precuneus. In a study of the elderly, males showed greater connectivity than females in the salience network, whereas females showed greater connectivity than males in the default mode network (Jamadar et al., 2018). Here, we demonstrate sex differences in somatomotor and occipital cortex, and cold color regions are the orbitofrontal cortex and temporo-parietal junction, suggesting that the ICA-CAP provides another analytical approach that may capture sex differences in network connectivity.

## 4 Discussion

In this study, we propose an approach that integrates group ICA and a covariance regression method. The covariance regression method identifies brain subnetworks that demonstrate population or individual variation in brain functional connectivity. Applied to the independent components (ICs) acquired from the group ICA, the proposed approach reconstructs principal component brain maps, comprised of orthogonal groups of the ICs, in association with covariates of interest. Compared to a standard pairwise approach, which requires fitting separate models for each pair of regions/networks, the utilization of the covariance regression method illustrates superior performance by avoiding the massive number of univariate tests.

Our method comparison adds a growing literature on comparing multivariate approaches to univariate approaches for functional connectivity modeling. The effectiveness of multivariate approaches over univariate ones was also observed for multivariate covariance measures (Geerligs et al., 2016; Yoo et al., 2019). In this first paper, we focused on multivariate modeling of whole correlation matrices, instead of each connectivity edge separately. The correlation matrices were computed using Pearson’s correlation, due to its wide adoption. However, this computation is considered to be “univariate”, compared with other “multivariate” correlations such as distant correlation (Székely et al., 2007). A future direction is to evaluate the potential improvement in reliability and validity by replacing Pearson’s correlation with distant correlation.

There are several methodological limitations of our current method. Given the current sample size, we didn’t consider functional connectivity changes over time or cognitive state, also known as dynamic connectivity (Hutchison et al., 2013). Additionally, spatial variations in functional connectivity were also found recently to be related to cognitive state (Salehi et al., 2019). Our framework takes a generalized linear model form, and this makes it amenable to inclusion of spatial and temporal covariates. Though estimating the spatial and temporal effects seems achievable, proper statistical inference would require future work to consider spatial and temporal dependence in a more complex mixed effects model framework. Secondly, Our method does not model task activations and even further task-induced connectivity changes. It marginally depends on the Gaussian distribution assumption, though Pearson’s correlation is relatively robust against slight departure of Gaussianity. Note that the non-Gaussian assumption over spatial maps in our group ICA does not apply to our Gaussian likelihood modeling of the extracted time courses. Future research is required to extend our method to accommodate various ICA approaches (Calhoun et al., 2009) with non-Gaussian assumptions on other components. Finally, we took a multi-stage approach which can lead to decreased statistical efficiency. An alternative approach, though computationally more expensive, is to consider fitting CAP regression and group ICA simultaneously in a uniform model. We apply the proposed method to the HCP resting-state fMRI data and identify brain subnetworks within which the functional connectivity variations can be explained by sex and/or alcohol use. Our findings are in line with extant literature, lending evidence to the usefulness of the proposed method in investigating the variability in brain connectomics. The main goal of the analysis herein is to assess the effectiveness of the proposed method. Future analyses with larger cohorts are warranted to validate the findings here. With increased cohort sizes and the availability of more comprehensive covariates, the propose method may be adopted to include more complex covariates and their interactions.

We also recognize several limitations in our fMRI analysis. First, we did not evaluate variations in brain maps related to covariates. Though these maps can be useful for generating hypotheses, our current implementation does not provide statistical significance of these maps or cluster-level *p*-values. One possible direction to use bootstrapped data to evaluate the variations in recovered brain maps, though this can be computationally prohibitive not to mention a potential challenge to match brain maps across bootstrapped samples. Second, it is expected that many other covariates could impact functional connectivity networks, for example structural imaging measures and behavioral assessments. In this first paper, we use the basic demographic variables in HCP as a demonstration of our method, and our conclusions are subject to confounding from those additional covariates not included in the model. Thirdly, we did not include additional data or external datasets to validate our findings. Here, our analysis should be treated as a confirmatory study illustrating a new method. The novel findings by our method should be further validated using ideally external data.

## A Additional Results of the Human Connectome Project Analysis

Figure A.1 is the boxplot of model outcome in the proposed ICA-CAP approach for each sex×alcohol subgroup. For example, in component C1 (Figure A.1a), we observe the difference between females and males among the alcohol users; as well as the difference between female alcohol drinkers and female non-drinkers. Figure A.2 presents the functional connectivity matrix of two subjects with the highest and lowest model outcome in CAP-C1. One subject (Subject 82) is female and a alcohol user, and the other one (Subject 86) is also female but claims no alcohol consumption. From the figure, we observe difference in connectivity patterns, where Subject 86 has strong functional connectivities in more IC pairs.

Using edge-wise regression on the two top loading ICs in components C1 and C3, Figure A.3 verifies the findings from the ICA-CAP approach. Figure A.4 shows the brains of components.

**Figure A.1:**
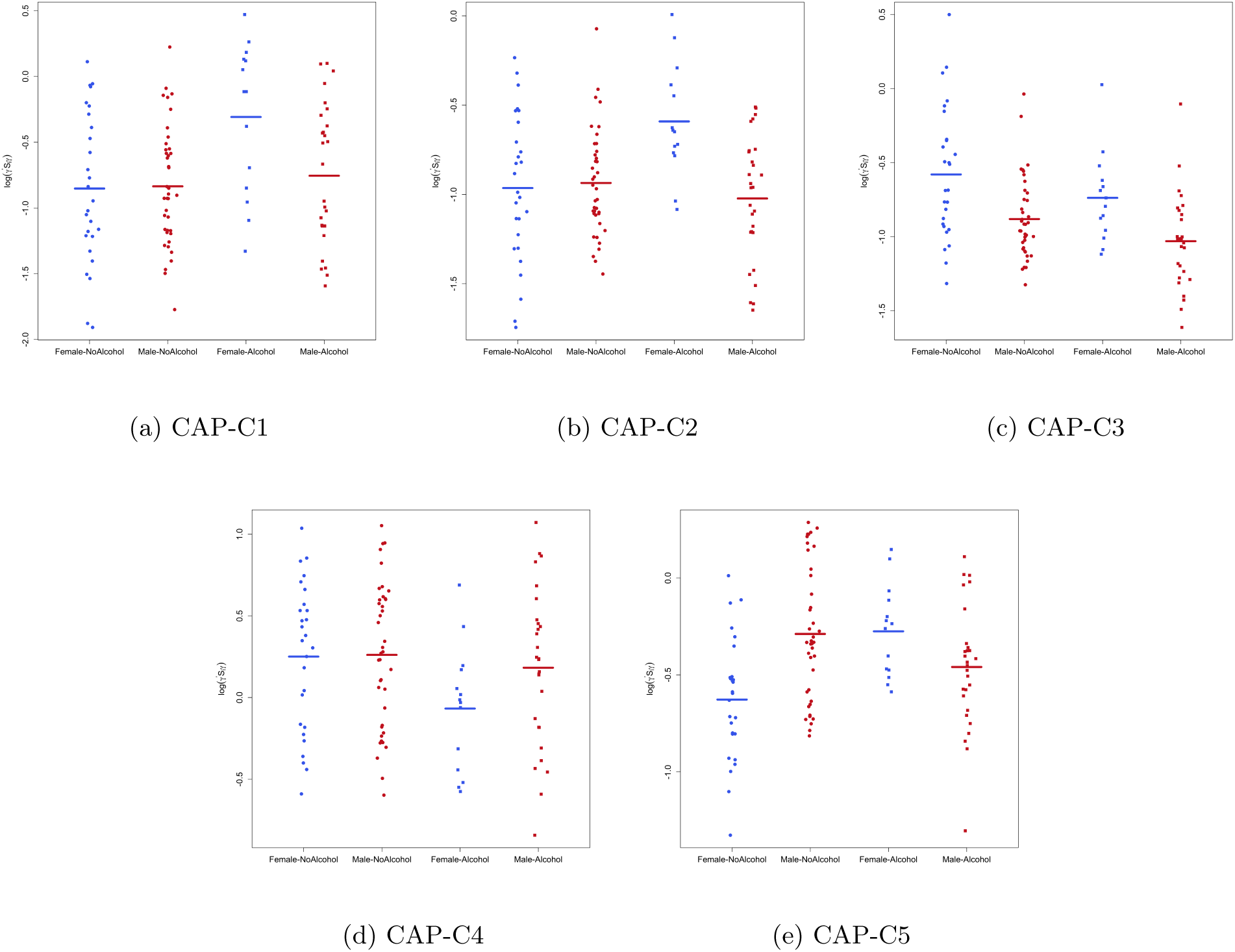
Scatterplot for the five identified components from CAP after adjusting for age. Female-NoAlcohol (blue solid circles): female alcohol non-drinkers. Male-NoAlcohol (red solid circles): male alcohol non-drinkers. Female-Alcohol (blue solid squares): female alcohol drinkers. Male-Alcohol (red solid squares): male alcohol drinkers.

**Figure A.2:**
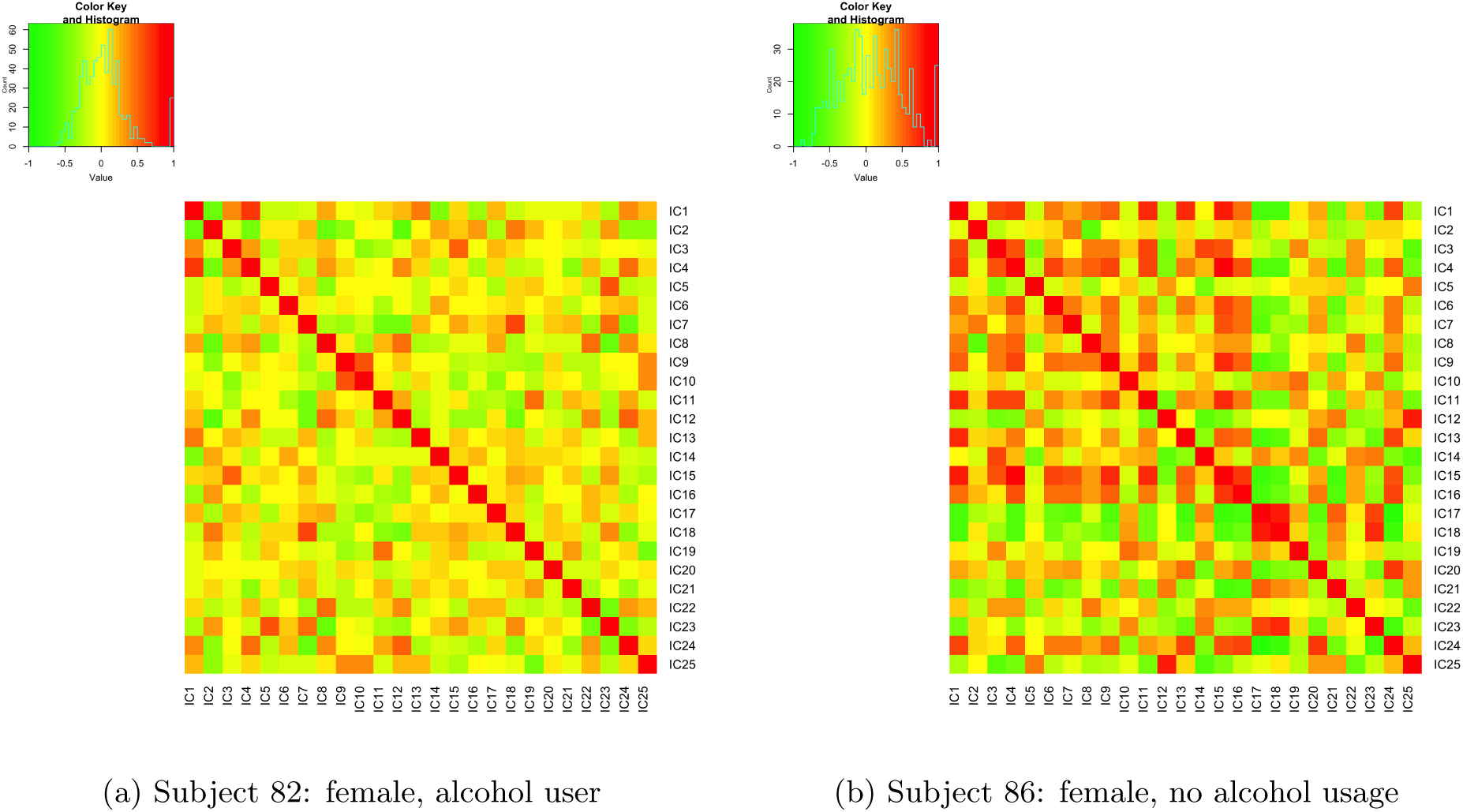
Functional connectivity matrix of two subjects with the highest and lowest model outcomes in CAP-C1.

**Figure A.3:**
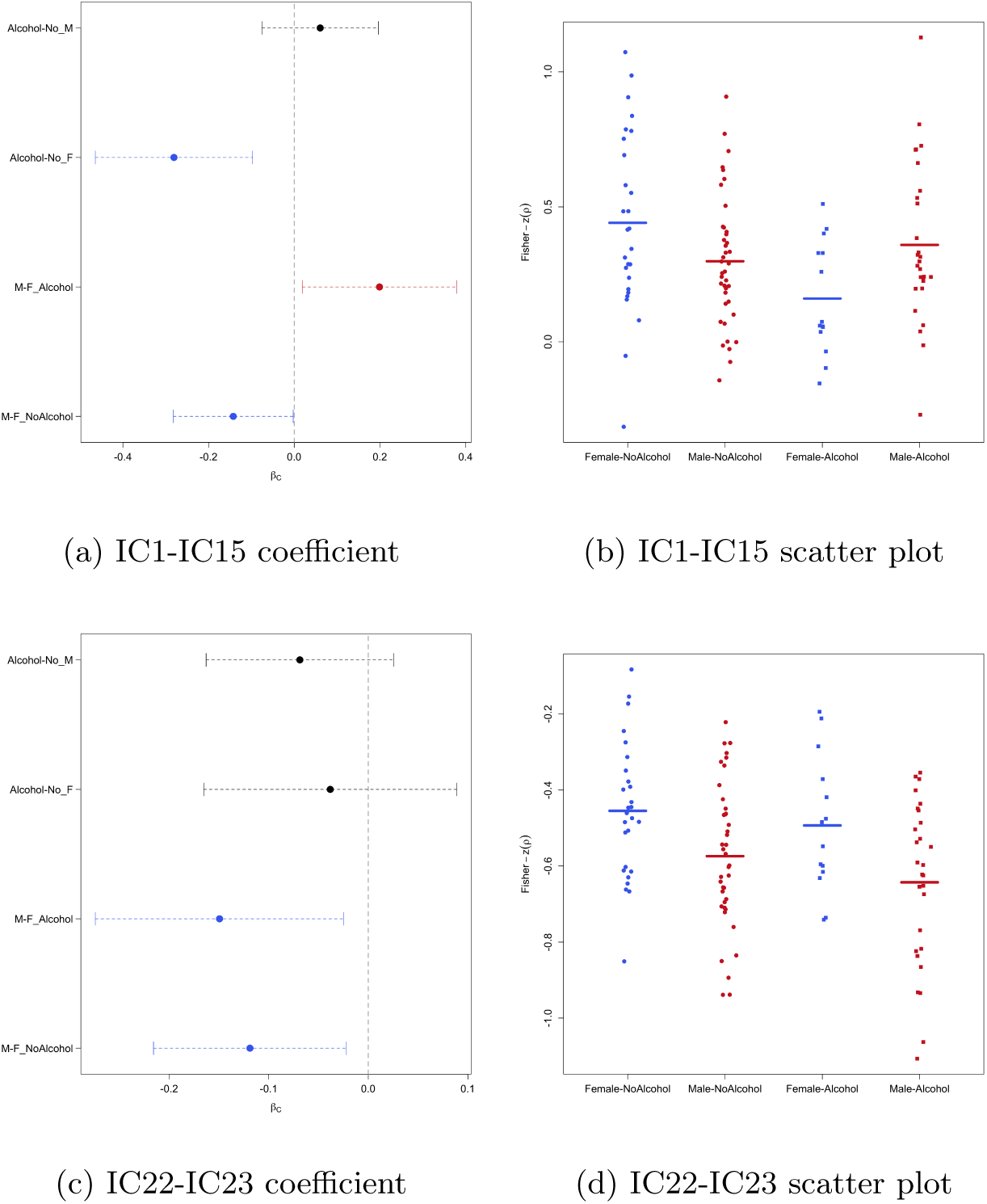
Estimated model contrasts of sex and alcohol (and 95% Bootstrap confidence intervals) and scatterplots (after adjusting for age) in the edge-wise regression for the top loading pairs in CAP-C1 and CAP-C3. In the scatterplot, Female-NoAlcohol (blue solid circles): female alcohol non-drinkers; Male-NoAlcohol (red solid circles): male alcohol non-drinkers; Female-Alcohol (blue solid squares): female alcohol drinkers; Male-Alcohol (red solid squares): male alcohol drinkers.

**Figure A.4:**
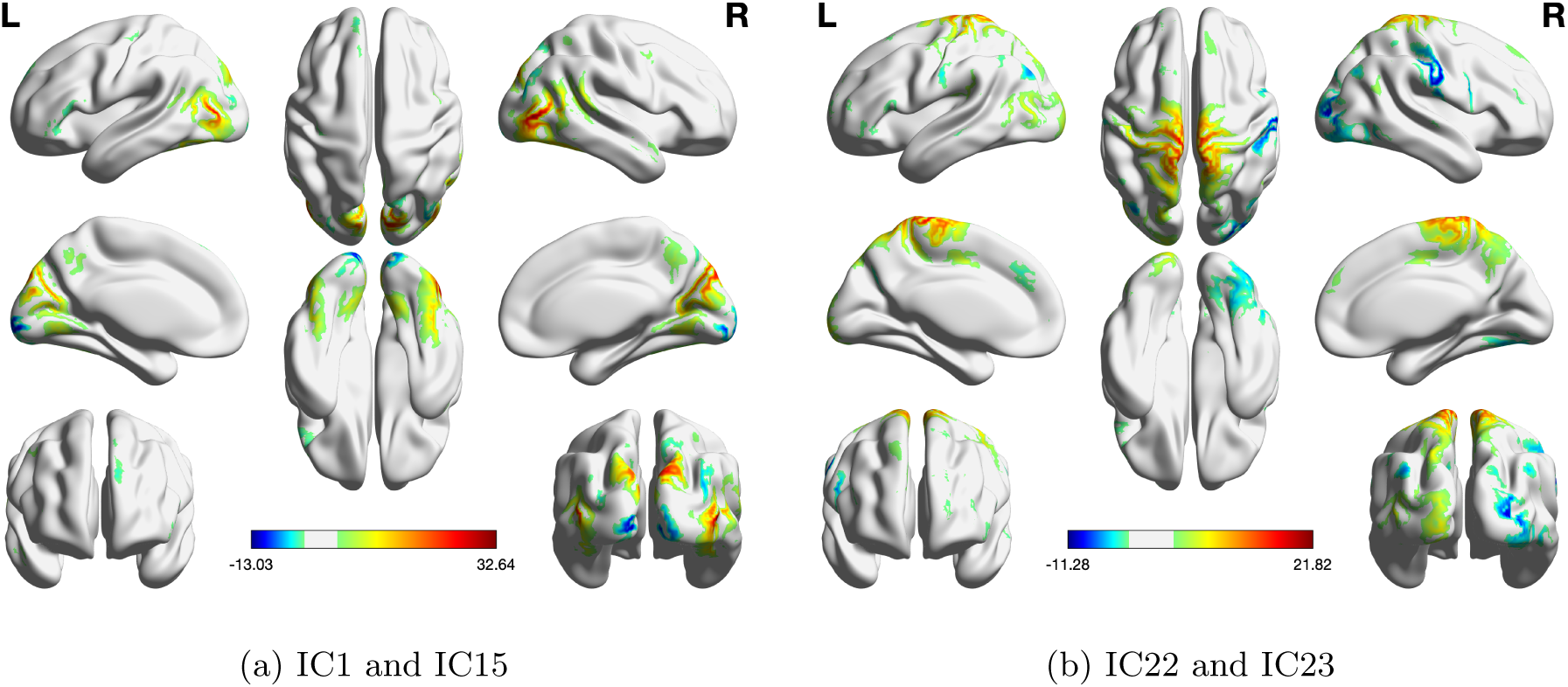
Brain maps of (a) IC1 and IC15 and (b) IC22 and IC23. In each brain map, the two ICs have the same weight.

